# The Ccm3-GckIII signaling axis regulates Rab11-dependent recycling to the apical compartment

**DOI:** 10.1101/2024.05.03.592387

**Authors:** Alondra S. Burguete, Yanjun Song, Amin S. Ghabrial

**Author notes:** co-first authors (Alondra S. Burguete, Yanjun Song).

## Abstract

Kinase cascades underlie many signaling pathways and are key regulators of development and morphogenesis. We have characterized a Hippo-like kinase cascade consisting of Thousand and One kinase (Tao), Germinal Center Kinase III (GckIII/Wheezy), and Tricornered (Trc) that plays an essential role in morphogenesis of tracheal terminal cell tubes in *Drosophila*. In this cascade, GckIII is the central kinase and is thought to act together with its binding partner, Cerebral Cavernous Malformations 3 (Ccm3). As suggested by its name, *Drosophila Ccm3* is the ortholog of a human vascular disease gene. As such, defining the Ccm3 pathway is critical to understanding both normal development and disease. Here we generate and characterize a null allele of *Ccm3* in *Drosophila*. We uncover a maternal contribution of *Ccm3* to embryonic development, show that maternal/zygotic null embryos have defective multicellular tracheal tubes, and that tracheal terminal cells derived from zygotic clones that also lack maternal *Ccm3*, show fully penetrant tube dilation defects identical to those we previously described for other pathway genes. We show that wildtype Ccm3 colocalizes with p-GckIII during early embryogenesis, and that in larval terminal cells, is found in the nucleus as well as associated with the apical membrane. We further demonstrate that *Mouse protein 25 (Mo25),* known to encode a protein that binds and stabilizes GckIII proteins in the active conformation, is required to prevent *Ccm3*-like tube dilations, showing that Mo25 and Ccm3, together with Tao, are required to fully activate GckIII, which directly phosphorylates and activates Tricornered (Trc). We show that this Ccm3 signaling cassette operates in other epithelial tissues such as the wing, and in non-epithelial tissues such as motor neurons. Lastly, we define a role of Ccm3-GckIII signaling in the distribution of active Rab11, leading us to propose that persistent local Rab11 activity results in elevated recycling of membrane and apical determinants to the apical domain, and consequent dilation of tubes. We validate this model by showing that loss of Rab11 activity ameliorates the tube dilation defects of pathway mutants.

## Introduction

The *Cerebral Cavernous Malformations 3* gene (*Ccm3*), was initially identified as encoding a protein upregulated during programed cell death (PDCD10) (Wang et al., 1999), but was subsequently found to be one of three genes whose loss of function can lead to familial cerebral cavernous malformations (Bergametti et al., 2005; Guclu et al., 2005). Familial CCM mutations are autosomal dominant, with variable expression and incomplete penetrance. Somatic loss of the remaining wild type copy (Knudsonian 2 hit mechanism) of a CCM gene is believed to result in localized vascular lesions (Pagenstecher et al., 2009) in which grossly dilated, thin-walled capillaries lacking surrounding support cells undergo repeated hemorrhage leading to headaches, neurological deficits and stroke (Haasdijk et al., 2012). Mutations in three genes were found to be sufficient to account for all or almost all cases of familial CCM. These genes – *Krev interaction trapped protein 1 (Krit1), Malcavernin,* and *Programmed Cell Death 10 (PDCD10)* – are commonly known as *CCM1*, *CCM2* and *CCM3,* respectively. The encoded CCM proteins are thought to act as scaffolding for regulators of cytoskeletal remodeling and/or signaling pathways; indeed, the proteins have been shown to physically interact to form a ternary complex (Hilder et al., 2007; Voss et al., 2007). Despite two decades of promising advances, and some early clinical studies, the state-of-the-art treatment for CCMs consist of palliative therapies and/or neurosurgical resection, when possible.

The precise functions of individual proteins in the CCM complex remain key subjects of inquiry. The current thinking in the field (Snellings et al., 2021) (Qi et al., 2023; Valentino et al., 2021) is that the CCM complex is required to negatively regulate Mitogen Activated Protein Kinase Kinase Kinase 3 (MAP3K3/MEKK3). The molecular mechanism by which MAP3K3 is regulated is unknown, although a physical interaction between CCM2 and MAP3K3 has been detected (Fisher et al., 2015; Wang et al., 2015). A recent study suggested bridging the kinase domain of GCKIII proteins with MAP3K3 may be key (Yang et al., 2023); however, no evidence of MAP3K3 phosphorylation by GckIII proteins has been described, and the CCM2-GckIII fusion protein may simply be a constitutively active GckIII isoform, leaving the mechanism of MAP3K3 regulation an unsolved mystery. Genetic evidence shows that reduction of MAP3K3 activity can decrease or prevent lesion formation in mouse CCM models (Zhou et al., 2016), whilst activating mutations in MAP3K3 have been found in the lesions of patients with sporadic CCMs (Hong et al., 2021; Weng et al., 2021). These data suggest that a critical function of the CCM complex is to negatively regulate MAP3K3 activity. Confounding the simplest models, however, loss of *MAP3K3* also causes brain hemorrhages (Deng et al., 2007), and vascular lesions associated with *MAP3K3* gain of function mutations seem to have a “popcorn-like,” appearance that is distinct from that of vascular lesions formed in familial CCM patients (Hong et al., 2021; Weng et al., 2021).

Here we focus on *Ccm3,* the *Drosophila* ortholog of human *CCM3*. Like *Ccm1* and *Ccm2*, post-natal endothelial-specific *Ccm3* knockout in mice results in formation of cavernous malformations; however, there are also data suggesting that vascular lesions (CCMs) result upon loss of *Ccm3* in neuroglia (Louvi et al., 2011), mural cells (Wang et al., 2020), or gut epithelia (Tang et al., 2019). In addition to the ternary complex that includes Ccm1 and Ccm2, Ccm3 is also found in other complexes, such as the STRIPAK complex that acts as an inhibitor of the Hippo pathway (Kean et al., 2011). Ccm3 has also been said to operate in a Ccm1- and Ccm2-independent manner even in endothelia (Chan et al., 2011; Yoruk et al., 2012; Zhu et al., 2010). Consistent with Ccm1/2-independent functions of Ccm3, global *Ccm3* knockout causes an earlier embryonic lethality in mice, and in humans, patients with *CCM3* mutations show a much earlier age of onset and a more severe course of disease (Chan et al., 2011; Denier et al., 2006; Shenkar et al., 2015). Likewise, patients with mutations in *CCM3*, but not *CCM1* or *CCM2*, develop multiple meningiomas at a high frequency (Riant et al., 2013). These differences hint at distinct downstream targets dependent specifically upon CCM3. Genetic data from zebrafish support this model, showing an additive phenotype upon *Ccm3* knockdown in *Ccm2* null embryos (Yoruk et al., 2012), consistent with independent functions of the CCM proteins in preventing cavernoma formation.

In our previous studies in *Drosophila*, we identified a requirement for Ccm3 pathway genes in preventing dilation of tracheal tubes (Antwi-Adjei, 2021; Hudson et al., 2023; Poon et al., 2018; Schweizer Burguete and Ghabrial, 2020; Song et al., 2013). In tracheal terminal cells homozygous mutant for components of the signaling pathway, focal tube dilations arose in correlation with higher levels of apical membrane proteins (Crumbs, aPKC, and p-Moesin), enrichment of apical Rab11, and ectopic localization of septate junction proteins (Song et al., 2013). Further, we were able to show that the pathway included an upstream kinase, Tao, that phosphorylates and activates GckIII, as well as a downstream kinase, Tricornered, that itself is phosphorylated and activated by GckIII (Poon et al., 2018). We have shown that Trc-interacting proteins Furry and Mob2 also act in the pathway in tracheal cells (Antwi-Adjei, 2021; Hudson et al., 2023). Importantly, a role for Ccm3-GckIII signaling in vertebrates has now been described in kidney, where loss of function has striking parallels to our work in trachea – including increased p-ERM staining and altered Rab11a distribution (Wang et al., 2021a). While flies have a single GckIII family member, vertebrates have at least 3 family members (*Stk24,25,26*) which complicates pathway analysis due to functional redundancy. Despite this challenge, *Stk24*/*Stk25* double knockout has recently been shown to induce CCM formation in mouse endothelium. Here we build on prior results, showing that Ccm3 and Mo25 are both required for GckIII activity, that pGckIII co-localizes with Ccm3, and that the alteration of Rab11 localization is specific for the active conformation of Rab11 and is functionally relevant.

## Results

### Knockout reveals maternal contribution of Ccm3 to embryonic multicellular tubes

To generate a null allele of *Ccm3,* we used a homologous recombination strategy. We designed our targeting construct such that it removes the entire *Ccm3* coding sequence (**Figure 1A**). Animals homozygous for the knockout died as first and second instar larvae. To examine terminal cells in third instar larvae, mitotic recombination was induced, and positively marked zygotic clones mutant for *Ccm3^ko^* were found to show a 61% penetrant terminal cell tube dilation defect (**Figure 1B,C**, n = 54). To test the possibility that a maternal contribution of *Ccm3* mRNA or protein was masking the *Ccm3* null phenotype, we generated females bearing germlines homozygous mutant for *Ccm3^ko^*. These females were crossed to males carrying either the *Ccm3^ko^* allele or a chromosomal deficiency – Df(3R)EXEL 6174) – that uncovers *Ccm3* and neighboring genes. Animals that lacked maternal and zygotic *Ccm3* died during embryogenesis with severe tracheal defects (**Figure 1D,E**). Embryos homozygous for *Ccm3^ko^*were indistinguishable from *Ccm3^ko^*/Df(3R)EXEL 6174 embryos. *Ccm3* maternal/zygotic null embryos showed dorsal trunk tubes that were elongated and irregular in diameter, with posterior regions of the tubes showing especially strong increases in tube diameter and sinuosity. Staining against GASP, a protein secreted into the embryonic tracheal lumen, suggests a possible trafficking defect, as levels of GASP in the lumen were reduced and cytoplasmic staining was detected at later embryonic stages than in wild type embryos (**Figure 1E-E”**). Staining against DE-Caderin revealed that the mutant cells of the dorsal trunk adapted rectangular rather than hexagonal shapes, and formed adherens junctions that appeared longer and more undulated than in control (**Figure 1 F,G,H**). In animals lacking maternal *Ccm3,* we induced mitotic recombination to generate *Ccm3^ko^* zygotic clones during early embryogenesis. These larvae were rescued to viability by zygotic expression of the paternal *Ccm3* allele, which is expressed in all cells except for the positively marked mutant zygotic clones. The *Ccm3* null cells – lacking both maternal and zygotic *Ccm3* – that developed as terminal cells showed fully penetrant (100%, n = 50) tube dilation defects like those we previously described for *Tao*, *wheezy/GckIII, fry* and *trc* (Antwi-Adjei, 2021; Hudson et al., 2023; Poon et al., 2018; Song et al., 2013). *Ccm3* null cells that formed autocellular tubes (stalk cells of the dorsal branch, for instance) also displayed focal tube dilations in which small sections of the autocellular tube bulged outwards into the surrounding cytoplasm of the cell (**Supplemental Figure 1)**. Combined, these results demonstrate that all three tracheal tube types (multicellular, autocellular and seamless **(Ghabrial et al., 2003)**) are affected by the loss of *Ccm3* function).

**Figure 1:**
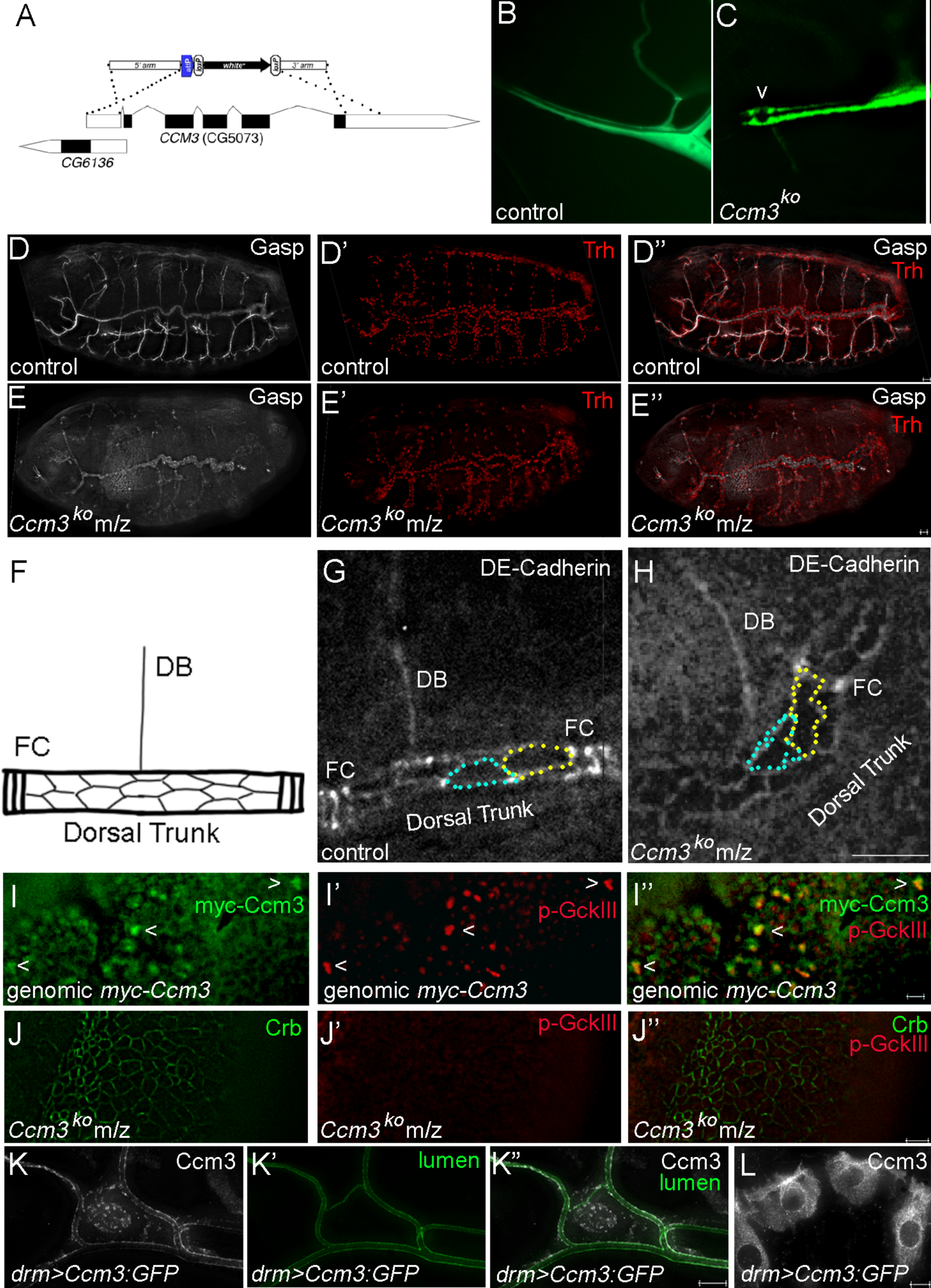
*Drosophila Ccm3* knockout causes tube dilation defects and eliminates p-GckIII staining. To analyze the requirement for *Ccm3* during *Drosophila* development we generated a *Ccm3* knockout strain using a targeting construct (A) and a homologous recombination strategy. The targeting construct was designed with homology arms flanking the coding sequence of *Ccm3*. A transgenic strain was generated and a targeting event was identified as described in (Huang et al., 2009). A single knockout line was recovered that removed all *Ccm3* coding sequence. In control terminal cells (B), a smoothly tapering tube extends from the intercellular junction with the neighboring stalk cell to roughly the position of the terminal cell nucleus, distal to which extensive branching of the primary tube occurs (one or two side branches are sometime found between the intercellular junction and the terminal cell nucleus). In *Ccm3^ko^* cells generated by mitotic recombination during embryonic stages, 60% of terminal cells showed tube dilation (C), indicated in this image by (<). To test whether the incomplete penetrance of the defect was due to maternal deposition of wild type *Ccm3* gene product, we generated females bearing germline clones null for *Ccm3* utilizing the female sterile technique (Chou et al., 1993). In zygotic *Ccm3* clones also lacking maternal *Ccm3*, terminal cells showed a 100% penetrant tube dilation defect (n=). In *Ccm3* maternal/zygotic null embryos, we observed tracheal tube defects in the multicellular dorsal trunk tubes. In control embryos (D, D’’), α-Gasp staining reveals the lumens of the tracheal tubular network. Compared to control, m/z *Ccm3* embryos (D, D’,E, E’) showed delayed Gasp deposition and lower signal intensity. Dorsal trunk tubes were found to be longer and more convoluted; the defects were particularly notable in the posterior segments of the embryos. The number of tracheal cells specified and contributing to specific tracheal tubes did not appear to be affected. Closer examination of dorsal trunk tubes looking at the apical surface area using antibody against the adherens junction component, DE-Cadherin, revealed that *Ccm3* mutant cells tended to have cell outlines that were more irregular, with the junctions appearing to be sinuous rather than straight, and to define rectangular rather than hexagonal shapes. The *Ccm3* null mutations were fully rescued by expression of a genomic myc-*Ccm3* construct in which a myc-tagged Ccm3 protein was expressed under the control of the endogenous *Ccm3* promoter (I). Staining against myc-Ccm3 in the early embryo revealed cell outlines and mitotic figures. We tested whether there might be co-localization between Myc-Ccm3 and GckIII phosphorylated on Threonine 167 (p-GckIII) (I), and found that there was. These findings suggest that the role of orthologous yeast proteins in mitotic exit might be conserved. In *Ccm3* m/z embryos, cells outlined with Crb staining (green, using a functional GFP knockin allele, (Huang et al., 2009)) lacked p-GckIII staining, consistent with Ccm3 being required for either activation of GckIII by Tao or another upstream kinase, or for stability of p-GckIII. Loss of *Ccm3* in tissue culture has been linked to activation and nuclear localization of YAP/TAZ (Wang et al., 2021b), however, we did not observe a change in Yki localization (cytoplasmic) in *Ccm3* m/z embryos (data not shown). To visualize Ccm3 subcellular localization in larval tracheal terminal cells, a UAS-Ccm3-GFP transgene was expressed using the *drum*-GAL4 driver (K). Ccm3-GFP was found to outline terminal cell tube lumens and to mark the nucleus; in contrast, in epidermal cells (L), Ccm3 was evenly distributed throughout the cytoplasm and was excluded from nuclei. Scale bars = 10 microns.

We tested whether expression of a wildtype *Ccm3* cDNA from a transgenic construct (UAS-*Ccm3-GFP*) would rescue zygotic tube dilation defects, and found that it did so (32/32 terminal cells with wild type appearance). Likewise, we showed that expression of a genomic rescue construct carrying an N-terminal Myc-tagged Ccm3 (genomic *myc-Ccm3*) and flanking genomic regulatory sequences, rescued animals that carried the *Ccm3^ko^* allele *in trans* to Df(3R)EXEL 6174; indeed, such animals are viable and fertile and have continuously propagated for multiple generations (the progeny would be a mix of *Ccm3^ko^* homozgyotes and *Ccm3^ko^*/Df hemizygotes, all carrying the rescuing *Myc:Ccm3* transgene). Together, these data demonstrate that our knock-out was specific for *Ccm3* and did not affect any other essential gene. Likewise, sequence data confirmed targeted deletion of the endogenous *Ccm3* locus in the knockout strain.

To verify that GckIII acts downstream of Ccm3, we tested whether *GckIII* transgenes could rescue the *Ccm3* terminal cell dilation defect. We found that overexpression of wild type GckIII lowered the penetrance of the *Ccm3^ko^* dilation defect to from 61% to 31% (n= 68), while expression of hyperactive GckIII (GckIII^HA^, that carries a T167D phospho-mimetic point mutation (Poon et al., 2018)) completely rescued *Ccm3^ko^*terminal cell tube dilations (100%, n = 50; we note that rescue was dosage sensitive, with the addition of another UAS transgene into the genetic background substantially reducing rescue). Another activated GckIII transgene provided incomplete rescuing activity (GckIII^DA^ – “doubly activated,” GckIII carrying the point mutation present in GckIII^HA^ (T167D) and truncated at the C-terminus to simulate potential caspase processing that has been proposed to activate some GckIII family members (Nogueira et al., 2008)).

### Ccm3 localization

To determine Ccm3 subcellular localization, we made use of the rescuing genomic *Myc-Ccm3* transgene. Myc-Ccm3 expression was detectable at early embryonic stages, but could not be distinguished from background in later embryonic and larval stages. In the early embryo (**Figure 1I**), Myc-Ccm3 was either cortical, or marked mitotic figures within a mitotic domain. Importantly, Ccm3 staining overlapped with p-GckIII (**Figure 1I’, I”**); further, in *Ccm3* maternal/zygotic embryos, pGckIII staining was not detected (**Figure 1J-J”**). To gain insight into Ccm3 localization in larval terminal cells, we used expression of GFP-tagged Ccm3. The *UAS-Ccm3-GFP* transgene was able to rescue *Ccm3^ko^*terminal cell tube defects, but GFP fluorescence was not readily detectable in the trachea by direct fluorescent microscopy. In *drum*-*GAL4*, *UAS-Ccm3-GFP* larvae, GFP signal could be detected in rows of epidermal cells but not in tracheal terminal cells; these GFP positive larvae were fixed and stained using an anti-GFP antibody to amplify the signal. Staining revealed that Ccm3-GFP was expressed, as expected, in tracheal terminal cells, and was enriched apically and in nuclei (**Figure 1K**). The pattern of localization was very reminiscent of that seen for p-Trc staining (Poon et al., 2018), consistent with Ccm3-GckIII activation of Trc by phosphorylation in these subcellular compartments (Poon et al., 2018). In contrast, in the higher expressing epidermal cells, Ccm3-GFP staining was strong, but was excluded from nuclei and was more broadly distributed throughout the cytoplasm in a punctate pattern (**Figure 1L**). These results raise the interesting question of whether there is a post-translational mechanism of Ccm3 turnover that limits Ccm3-GckIII activity.

### Mo25 is part of the Ccm3 signaling cassette and acts upstream of GckIII

Yeast do not have a Ccm3 ortholog, but do have GckIII proteins that phosphorylate Trc orthologs (Nelson et al., 2003). To generate fully active their GckIII kinases, yeast require a Mouse protein 25 (Mo25) ortholog (Hsu and Weiss, 2013). These conserved signaling pathway components function in the so-called Mitotic Exit Network (MEN) and the Regulation of Ace2 activity and cellular Morphogenesis (RAM) network (Duhart and Raftery, 2020). In *Drosophila*, Mo25 is conserved, but was reported to be dispensable for Trc function, suggesting that the pathway may differ from yeast (He et al., 2005); however, in our hands knockdown of *Mo25* appeared to affect Ccm3 pathway activity in the trachea (Hudson et al., 2023), consistent with findings in *c. elegans* (Lant et al., 2018). To definitively test the potential function of *Mo25* in trachea, we initially characterized the *Mo25^Δ8-2^* P-element excision allele (lost during Covid shut down), and subsequently validated our observations with newly generated Crispr alleles of *Mo25*. We examined mosaic animals and found that *Mo25* deficient terminal cells showed a 100% penetrant tube defect (n = 50) indistinguishable from that of *Ccm3* or *wheezy/GckIII* null cells (**Figure 2A,B**). Expression of *UAS-Mo25* was able to rescue *Mo25* loss of function (**Figure 2C**), but not *Ccm3^ko^* or *GckIII^770^* (**Figure 2D,E**).

**Figure 2.**
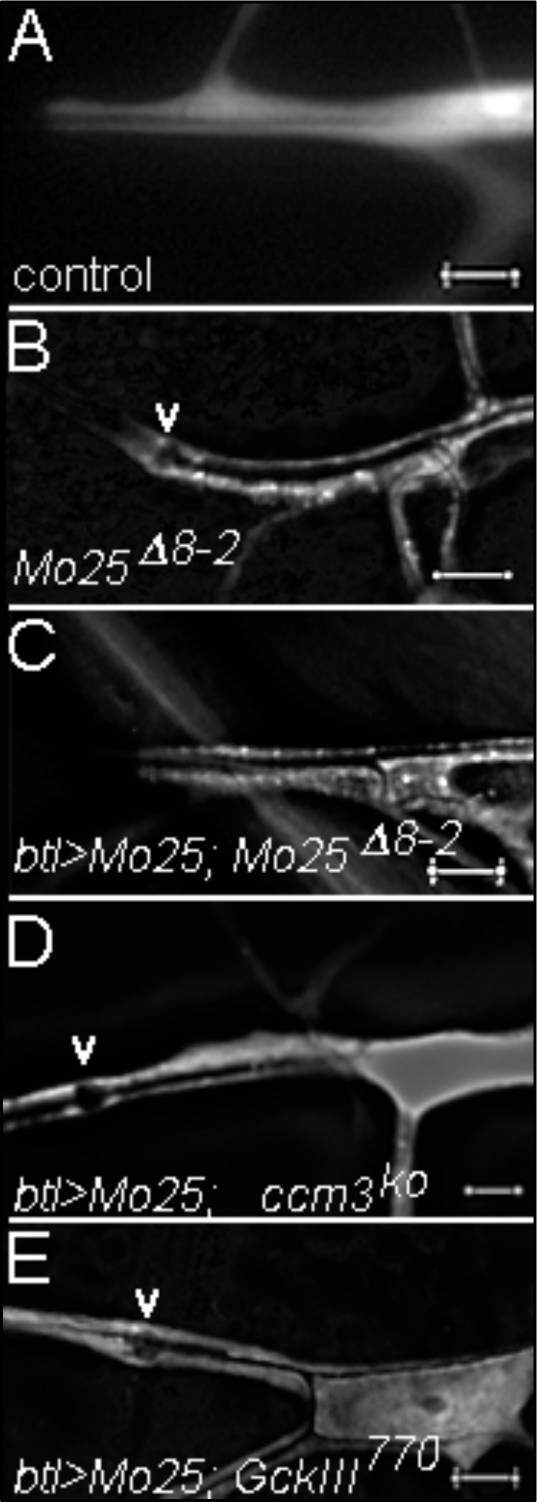
Epistatic relationships among *Mo25*, *Ccm3* and *GckIII*. In control (A) terminal cell clones (positively labeled with GFP, white), a single tube smoothly tapers as it runs from the intercellular junction into the cell soma where it will branch (some cells will branch once or more prior to reaching the terminal cell nucleus). In *Mo25^Δ8-2^* terminal cell clones (B), one or more large focal dilations (marked with “v”) are detected between the intercellular junction and the cell soma. In *Mo25^Δ8-2^* terminal cells expressing wild type Mo25 from a transgene (C), no tube defects are observed. Over-expression of wild type Mo25 cannot suppress tube dilations (“v”) caused by mutations in Ccm3 (D) or GckIII (E). Scale bars = 10 microns.

GckIII^HA^ expression was sufficient to bypass the requirement for *Mo25* and *Ccm3* in terminal cells (n = 50, n = 50, respectively). Intriguingly, and in contrast to the case for *Ccm3*, overexpression of wild type GckIII did not show any rescuing activity (data not shown) for *Mo25* mutant cells. These data are consistent with biochemical work showing that Mo25 promotes the active GckIII conformation (Filippi et al., 2011), and that Ccm3 binds and stabilizes GckIII (Fidalgo et al., 2010). It remains to be determined whether Ccm3 and Mo25 physically interact, or if Mo25 facilitates Ccm3 binding of GckIII. In *C. elegans*, Mo25 has been reported to be essential for Ccm3 localization; however, whether this is direct or reflects Mo25 interaction GckIII is unclear (Lant et al., 2018).

### Ccm3 functions in the planar cell polarity/wing hair morphogenesis pathway

When we generated females with *Ccm3^ko^* germline clones, we also noted the presence of somatic clones in the wing. Wings bearing clones appeared irregular, with a wavy surface. Upon closer examination, unmarked *Ccm3^ko^* clones were detectable, showing a multiple wing hair defect similar to *trc* (Geng et al., 2000). We verified that the hair defect in mosaic animals reflects the loss of *Ccm3* in wing epithelia by utilizing *nubGAL4* to induce tissue-specific knockout by Crispr/gRNA expression, and also by tissue specific knockdown with *Ccm3* RNAi. Animals mosaic for *Mo25^Δ2^* also showed wing hair defects, and tissue-specific knockout of *Mo25* confirmed the role of *Mo25* in wing hair morphogenesis, contrary to published results (He et al., 2005). Compared to wild type wings (**Figure 3A**), clones homozygous for *Ccm3^ko^*(**Figure 3B**) or for *Mo25^Δ2^* (**Figure 3C**) showed *trc*-like multiple wing hair defects. We also tested for genetic interactions with *disheveled (dsh)*, a core component of the planar cell polarity signaling pathway (Axelrod, 2001). In *dsh^1^* mutant males, wing hairs are disoriented, and some cells produce more than 1 wing hair (1.05 wing hairs/cell +/- .01 SEM). In *dsh^1^* mutant males heterozygous for *GckIII^770^*, a significant increase was observed in the number of wing hairs per cell (1.35 +/- .03 SEM; p = 0.0001). In a control outcross, *GckIII^770^* heterozygous flies (wild type for *dsh*) had 1 wing hair (1 +/- 0.0 SEM).

**Figure 3:**
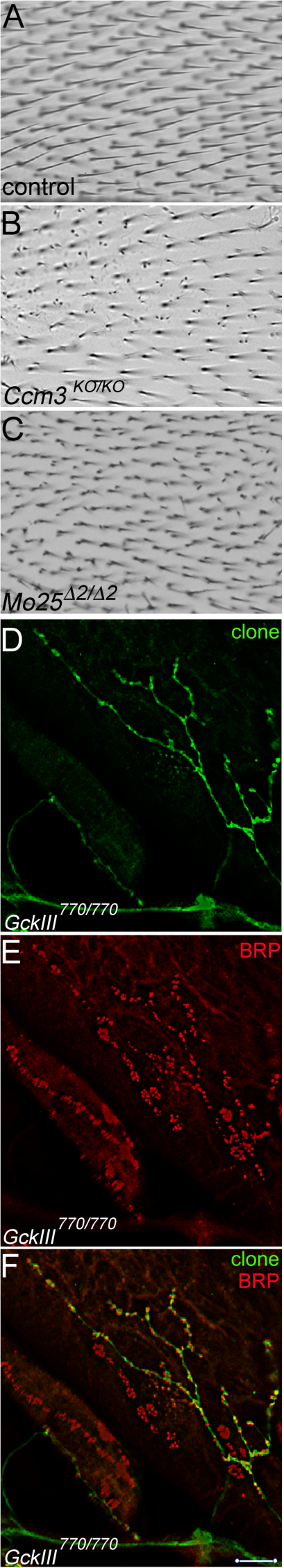
The Ccm3 signaling cassette operates in other epithelial tissues and neurons. In control wings, actin-based projections form single wing hairs at the posterior margin of each cell. The wing hairs extend distally (A). The Planar Cell Polarity genetic network detects and propagates polarity within the plane of the epithelium, allowing cells to coordinate their orientation across the epithelial tissue (Peng and Axelrod, 2012; Strutt and Strutt, 2021). PCP component Multiple Wing Hairs (MWH) is also required to ensure that single hairs are formed at the correct position in every cell. Cells mutant for *mwh*, or for *trc* or *fry* produce multiple hairs (Fang and Adler, 2010; Yan et al., 2008). We find that cells mutant for *Ccm3* (B) or *Mo25* (C) likewise form multiple wing hairs. In neurons, *trc* has been found to affect morphology of dendrites and synapses ((Emoto et al., 2004; Natarajan et al., 2015)). In motor neurons, mutations in Ccm3 pathway genes *Tao*, *trc*, and *Mob2* have been shown to cause the neuromuscular junction to form more boutons of smaller size (Campbell and Ganetzky, 2013; Natarajan et al., 2015; Politano et al., 2019). We find that motor neurons mutant for *GckIII* (D-F) or *Ccm3* display similar defects. Homozygous mutant motor neuron is labeled with GFP (green), while NMJ active zones of each bouton (mutant and wild type) are labeled by Bruchpilot (Brp, red). Scale bar = 10 microns.

### Ccm3 pathway is utilized in other tissues including neurons

Having found that the Ccm3-GckIII pathway regulates Trc in wing as well as trachea, we wanted to determine whether the signaling cassette acts in non-epithelial cells with a known *trc* requirement. We therefore turned to motor neurons, where other groups have identified requirements for *Tao*, *trc* and *Mob2* in bouton morphology (Campbell and Ganetzky, 2013; Natarajan et al., 2015; Politano et al., 2019). We found that *Ccm3* and *GckIII* loss of function altered neuromuscular junctions in a similar manner, resulting in more and smaller boutons (**Figure 3D-F**), compared to neuromuscular junctions formed by neighboring heterozygous neurons.

### The Ccm3 pathway regulates local activity of Rab11-GTP

Downstream targets of the Ccm3 pathway kinase cascade remain elusive. Based on our analysis of the cellular defects associated with loss of pathway function in trachea – which include tube dilation, accumulation of elevated levels of apical determinants and polarity proteins (Crumbs, aPKC, p-Moesin), altered GASP localization in m/z embryos, and elevated Rab11 staining in proximity to tube dilations in *GckIII* mutant terminal cells – we hypothesized that regulators of protein trafficking could be targeted. We examined the localization of YFP-Rab11 isoforms (WT, DN [S29N], and CA [Q77L]) in wild type and mutant backgrounds. We found that in a wildtype background, all three Rab11 isoforms were evenly distributed throughout the terminal cell cytoplasm (**Figure 4A** and data not shown). However, in *GckIII* mutant terminal cells, YFP-Rab11CA was uniquely enriched at sites of tube dilation and was overall more apically enriched and uneven in its distribution. We tested YFP-Rab11CA distribution in *Ccm3^ko^*zygotic clones (maternal *Ccm3* **not** eliminated) and found a similar but less robust change in YFP-Rab11CA localization (**Figure 4B, C**). Strengthening this connection to Rab11 localization, recently published data shows altered Rab11a distribution in Ccm3 pathway mutations in mouse kidney (Sartages et al., 2022). To determine if this alteration in Rab11 distribution merely serves as a corelative marker of pathway dysfunction, or if Rab11 is an important downstream effector, we tested whether mutations in *Rab11* could modulate the tube dilation defect of *GckIII^770^* terminal cells. We first characterized the *Rab11* loss of function phenotype in larval terminal cells. Tracheal terminal cells mutant for *Rab11^j2D1^*(**Figure 4D**) displayed partial gas-filling defects with more distal tubes lacking gas. In cells doubly mutant for *GckIII^770^* and *Rab11^j2D1^* (100% no dilations, n = 17; **Figure 4E**), or mutant for *GckIII* and expressing *Rab11^DN^* (98% no dilation, n = 45; **Figure 4F**), we found that the *GckIII* tube dilation defect was suppressed. These data imply that altered localization of active Rab11 is functionally relevant to the tube dilation defects observed upon loss of Ccm3 pathway function.

**Figure 4:**
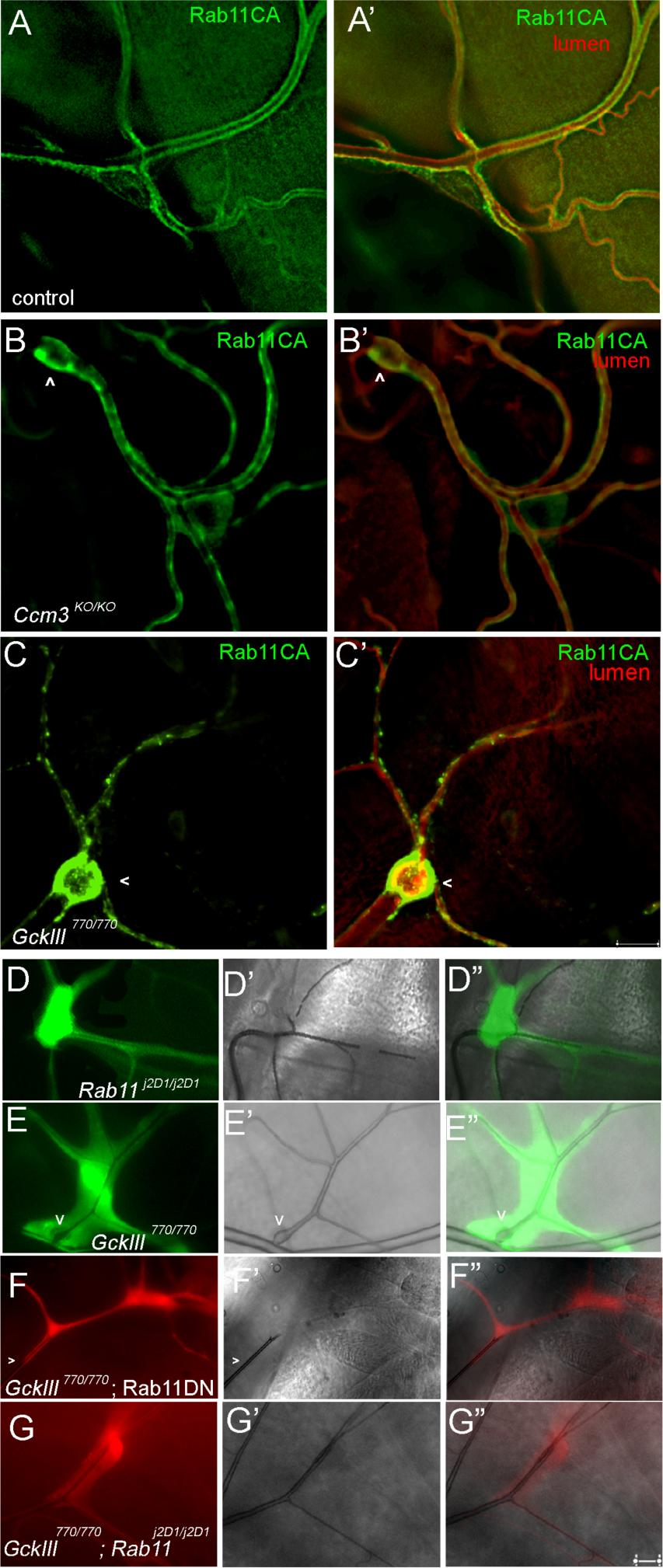
Rab11 localization and activity are altered in Ccm3 pathway mutants. Expression of YPF-Rab11 isoforms (WT, CA, and DN) was analyzed in wildtype and mutant backgrounds. We found that in a wildtype background, (A) all isoforms were uniformly distributed in terminal cell cytoplasm, but excluded from the nucleus. In *Ccm3* (B) and *GckIII* (C) zygotic loss of function clones, terminal cells showed patchy apical enrichment of Rab11CA but not Rab11WT or Rab11DN. In A-C, YFP staining (green) and lumenal staining (red) are shown. Clones were marked with RFP (not shown) using the MARCM system. Altered distribution of Rab11CA was less pronounced in *Ccm3^KO^* (maternal *Ccm3* is present) than in *GckIII* mutant cells. In *Rab11^J2D1^* mutant terminal cells (D) a partially penetrant gas-filling defect is seen, with more distal branches lacking gas-filling. The image shown is a composite of 4 different Z sections of the same cell. In *GckIII^770^*mutant terminal cells (E) a large transition zone dilation is detected (marked by v). In terminal cells mutant for *GckIII^770^* and also expressing a dominant negative Rab11 transgene (F) or doubly mutant for *GckIII^770^*and *Rab11^J2D1^* (G), transition zone tube dilation is suppressed.

## Discussion

The Ccm3 Hippo-like signaling pathway is required to prevent tube morphogenesis defects in *Drosophila* tracheal tubes, *c. elegans* excretory cell tubes, and vertebrate cerebral endothelial tubes. Our published data shows that the Ccm3 pathway includes an upstream activating kinase, Tao, that can promote both Hippo pathway and Ccm3 pathway signaling, begging the question of whether the two pathways are always activated in tandem, or whether other factors can promote selective activation of one pathway over the other. Tao phosphorylates and activates GckIII, but full GckIII activity also requires the scaffold protein Ccm3, that might function to localize or stabilize the kinase, as well as Mo25, whose binding is thought to promote the active conformation of GckIII. In turn, GckIII phosphorylates and activates Trc. Full activity of Trc also requires a scaffolding protein, Furry, and an activator, Mob2. The precise mechanisms by which Furry promotes Trc activity are not clear, and whether Mob2 is the sole Mob family member required to regulate the Ccm3 pathway is likewise uncertain. Physical interactions between Trc and both Mob1/Mats and Mob2 have been described (Duhart and Raftery, 2020). However, in the trachea, loss of function of the Hippo pathway and of the Ccm3 pathway are phenotypically distinct, suggesting little redundancy between Mob1 and Mob2. Very little is known about Mob3, but Mob4 is a known component of the STRIPAK complex that is thought to negatively regulate Hippo signaling, and that also contains Ccm3 and GckIII proteins (Goudreault et al., 2009).

Prior work in embryos suggested that Rab11 vesicles prefigure seamless tubes, and would therefore contribute to their growth (Gervais and Casanova, 2010). Other data show larval seamless tubes also depend on endocytic trafficking (Jones et al., 2014; Schottenfeld-Roames et al., 2014). Combined, these observations and our own suggest that Rab11 regulation of endocytic recycling plays a key role in lumen morphogenesis. How the Ccm3 pathway regulates Rab11 activity remains a key question. In *c. elegans*, loss of *Ccm3* function is thought to compromise endocytic trafficking in the excretory canal cell – a single celled tube – and the multicellular germ cell tube, through a CDC42-dependent mechanism (Lant et al., 2018; Lant et al., 2015; Pal et al., 2017). We do not yet have data to support a CDC42-dependent function of Ccm3 in *Drosophila*. In principle, Rab11 regulation could be direct; indeed, mass spectrometry analysis has revealed that Ser 177 of Rab11 is phosphorylated *in vivo* and that the phosphomimetic mutant, Rab11 S177D, displays altered transferrin trafficking (Pavarotti et al., 2012). To date, we have not detected any change in Rab11 phosphorylation in *Ccm3* pathway mutants. Alternatively, less direct mechanisms of Rab11 regulation could be at play, like targeting of Rab11 effectors such as the *Drosophila* Rab11FIPs, Nuf and Rip11. Other potential targets could include Rab11GEFs or GAPs. Intriguingly, the yeast and mammalian orthologs of Trc kinase have been shown to phosphorylate client proteins on a consensus sequence of **H**X**R(R/X**)X(**S*/T\***), with a strong preference for H in the -5 position (Mazanka et al., 2008; Ultanir et al., 2012). Importantly, Rab11FIP5 (orthologous to Rip11) has been identified in a chemical genetic screen as a Stk38/38l (NDR1/2) client protein, and in mouse *Stk38/38l* double knockout neurons, mass spectrometry showed that Rab11FIP5 had a dramatic loss of the corresponding phosphosite (Rosianu et al., 2023; Ultanir et al., 2012). Conservation between fly Rip11 and mouse Rab11FIP5 does not include the identified phosphosite, however, HXXXXS motifs are present in fly Rip11, so phospho-regulation of Rip11 could be functionally conserved.

**SUPPLEMENTAL FIGURE 1:**
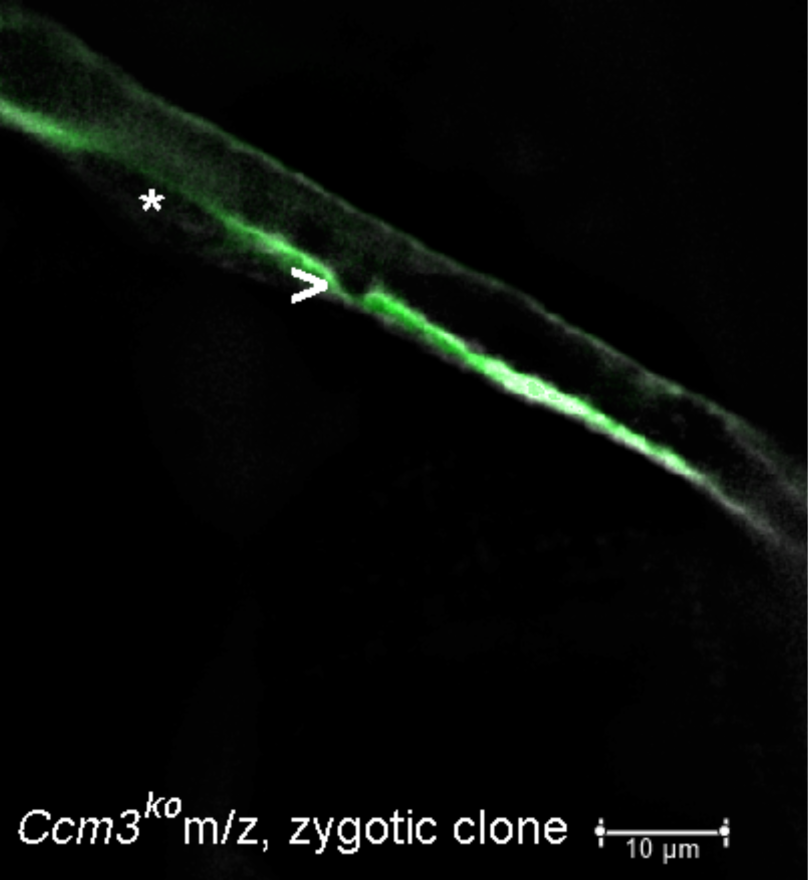
Autocellular tubes deficient for Ccm3 also show focal dilations. In this autocellular tube, a single cell is wrapped around a lumenal space and forms a self-junction running along its long axis. This particular cell is part of a zygotic clone homozygous mutant for *Ccm3^ko^*. The mosaic animal in which the clone was induced also lacked maternal *Ccm3*. Note the small focal dilation in the tube (and outward bulging indicated by the >). The position of the cell’s nucleus is indicated by *. Scale bar = 10 microns.

## MATERIALS AND METHODS

### Construction of *Ccm3^ko^*

The *Ccm3^ko^* allele was generated according to the protocol established by the Hong lab ((Huang et al., 2009)). *Ccm3* 5’ and 3’ homology arms were amplified by PCR as follows:

5’ homology arm (2.935 kb):

5’-ATAAGAATGCGGCCGCCCTTGGATGCGTTTTCTTC-3’

5’-CTAGCTAGCCACAACAGCACAGGCACAGT-3’

PCR was with Phusion high fidelity DNA polymerase (NEB). PCR product was TA tailed with TAQ and TopoTA cloned into pCR4. Subcloning into pGX-attP was as a Not I, Nhe I fragment. 3’ homology arm (5.019 kb):

5’-GAAGATCTATACAAAATCGCCCGACGCAGTCC -3’

5’-AACTGCAGCTCCACACAACCACACATACG-3’

PCR and TA cloning as above. Sucloning into pGX-attP was as a Bgl II, Pst I fragment. The knockout allele was generated on a third chromosome carrying FRT82B to permit mosaic analysis.

### Mo25 Crispr alleles

Bloomington stocks (gRNA and nos-CRISPR) were utilized to generate small deletions in the Mo25 gene on an FRT^2A^ chromosome.

### Myc:Ccm3 transgenic strain

### UASt transgenic strains

*Ccm3:GFP* was constructed with Ccm3 amplified from cDNA clone RE18871 and EGFP amplified from pEGFP-N1*. GckIII^DA^* was created by the addition of a stop codon to the previously described GckIII^CA^ construct. The *Mo25* was made using a cDNA template and cloned as a XhoI - EcoRI fragment into pUASt.

### Wing hair analysis

wings were dissected from young adults, washed in 70% EtOH and mounted in Euparal. For counting of wing hairs per cell, ∼ 50 cells were scored, per wing, in a designated square region of interest posterior to the humeral crossvein. The p values were determined by unpaired T test using Graph Pad to analyze the data.

### Generation of zygotic clones by mitotic recombination

To evaluate the cell autonomous function of essential genes, a common approach is to generate clones of homozygous mutant cells in heterozygous animals. Our system for doing so has been described (Ghabrial and Krasnow, 2006), and permits positive labeling of mutant cells with GFP together with labeling of all cells in the tissue of interest with DsRed or mKate.

### Generation of *Ccm3^ko^* germline clones and m/z null embryos

We used the female-sterile technique (Chou et al., 1993) to generate females carrying germlines homozygous mutant for the *Ccm3^ko^* allele. These females were obtained by taking males from Bloomington stock 2149 (FRT82B p{ovoD1-18} and crossing them to virgins of the genotype: y, w, hsFLP122; Sp/CyO; MKRS/TM2, Gmr-Hid. The y, w, hsFLP122; FRT82B Ccm3ko/TM2,Gmr-Hid or MKRS male progeny were selected and crossed to virgins of the genotype: btl>wkdmkate/CyO; FRT82B *Ccm3^ko^*/TM3 Sb, Twi>EGFP. Larvae were subjected to daily heatshocks of 1 hour at 38.5 °C. Germline mosaic females were selected based on the absence of a 3^rd^ chromosome balancer, and were fertile when crossed to wild type males. To generate maternal/zygotic null embryos, the germline clone bearing females were crossed to males carrying either *Ccm3^ko^* or Df(3R)EXEL 6174 (Bloomington stock 7653) in *trans* to a “green” balancer TM3 Sb, Twi>EGFP (Bloomington 6663).

### p-GckIII antibody

(Genscript). The phosphopeptide SKRN(pT)FVGTPFC was synthesized and New Zealand white rabbits were injected. For Rabbit 5910, the unphosphorylated SKRNTFVGTPFC peptide was used to adsorb all non-phosopho-specific antibodies and the remaining sera was affinity purified.

